# Video-Based Cattle Behavior Detection for Digital Twin Development in Precision Dairy Systems

**DOI:** 10.1101/2025.11.13.688203

**Authors:** Shreya Rao, Eduardo Garcia, Suresh Neethirajan

## Abstract

Digital twins in dairy systems require reliable behavioral inputs. We develop a video-based framework that detects and tracks individual cows and classifies seven behaviors under commercial barn conditions. From 4,964 annotated clips, expanded to 9,600 through targeted augmentation, we couple YOLOv11 detection with ByteTrack for identity persistence and evaluate SlowFast versus TimeSformer for behavior recognition. TimeSformer achieved 85.0% overall accuracy (macro-F1 0.84) and real-time throughput of 22.6 fps on RTX A100 hardware. Attention visualizations concentrated on anatomically relevant regions (head/muzzle for feeding and drinking; torso/limbs for postures), supporting biological interpretability. Structured outputs (cow ID, start-end times, durations, confidence) enable downstream use in nutritional modeling and 3D digital-twin visualization. The pipeline delivers continuous, per-animal activity streams suitable for individualized nutrition, predictive health, and automated management, providing a practical behavioral layer for scalable dairy digital twins.

## 1. Introduction

Digital twin technology represents a paradigm shift in precision livestock farming, transforming how dairy operations monitor, analyze, and optimize animal management through the creation of virtual replicas of physical farm entities ^1, 2^. These advanced systems integrate real-time sensor data, behavioral patterns, physiological parameters, and environmental conditions to enable continuous monitoring, predictive analytics, and automated decision-making at scales and levels of precision previously unattainable ^3,4^. In the dairy sector, digital twins combine behavioral monitoring, mechanistic physiological modeling, nutritional requirement assessment, and environmental impact analysis to generate individualized management strategies that improve both animal welfare and production efficiency ^5^. Empirical evidence supports these benefits: digital twin deployment can enhance feed conversion efficiency by 15-20%, reduce veterinary interventions by 25-30%, and cut greenhouse gas emissions per unit of milk by as much as 25% ^6,2^. These outcomes illustrate the potential of digital twins as transformative tools for addressing the economic, environmental, and welfare challenges facing modern dairy production. Robust and continuous behavioral monitoring thus serves as the foundation upon which these digital twins operate, providing dynamic, high-frequency data to model individual cow states and management outcomes.

A central prerequisite for digital twin functionality is accurate and continuous behavioral monitoring. In dairy cattle, five behavioral categories like feeding, drinking, lying, standing, and ruminating serve as critical indicators of metabolic balance, reproductive status, disease onset, and welfare status ^7^. Feeding behavior is closely tied to dry matter intake and energy balance, and its measurement is essential for nutritional modeling and ration optimization. Rumination patterns provide direct insights into digestive function and metabolic efficiency, with deviations often preceding metabolic disorders such as acidosis, ketosis, or displaced abomasum ^5^. Lying behavior is a proxy for cow comfort and lameness risk; research has established that 10-14 hours of daily lying is optimal for both productivity and welfare outcomes^8^. Drinking behavior, while less frequent, is vital for assessing hydration, feed palatability, and heat stress, all of which impact milk yield and welfare^9^. Even standing, often overlooked as a passive activity, can indicate restlessness, discomfort, or estrus expression when measured systematically. Automated recognition of these behaviors is therefore indispensable for creating the continuous, structured data streams that feed into digital twin models ^1,3^. While the biological relevance of these behavioral metrics is well established, the challenge lies in capturing them continuously and objectively in commercial farm environments.

Conventional behavioral monitoring methods, however, present significant limitations that restrict their scalability and accuracy. Manual observation, though capable of capturing fine-grained detail, is labor-intensive, subjective, and economically unsustainable for commercial herds that often number in the hundreds. Even among trained observers, inter-observer variability can reach 15-25%, reducing the reliability of datasets essential for predictive modeling ^7,10^. Wearable sensor systems, while offering continuous monitoring, face a series of practical and welfare-related challenges. Device loss rates of 5-15% create gaps in longitudinal tracking. Batteries require replacement every 3-6 months, imposing labor costs and necessitating animal handling that can induce stress. Calibration drift reduces measurement reliability, while sensor placement may alter natural behaviors, potentially biasing the very data intended for welfare and productivity optimization ^5,6^. These shortcomings underscore the need for alternative approaches capable of generating high-quality, scalable behavioral data.

Computer vision has emerged as the most promising solution to these limitations, offering a scalable, non-invasive method for monitoring livestock at both individual and group levels. Barn-based video systems enable continuous monitoring of multiple animals simultaneously without interfering with their natural behaviors, while providing richer postural and temporal information than wearable sensors. Video analytics can capture body orientation, head movements, and activity transitions that are critical for distinguishing behaviors such as feeding versus ruminating. Importantly, video-based systems scale cost-effectively, monitoring dozens of animals with a single camera, and integrate easily with existing farm surveillance infrastructure (^11, 2^). Furthermore, standardized output formats from computer vision pipelines facilitate integration with farm management software, nutritional modeling tools, and digital twin frameworks.

Recent advances in deep learning architectures have dramatically improved the accuracy and robustness of behavior recognition systems. State-of-the-art models have achieved 85-95% accuracy in multi-class livestock behavior classification tasks ^3,10,12^. Transformer-based models, originally developed for natural language processing and now adapted for video understanding, show promise for long-duration behaviors. For example, transformer-based classifiers have reached over 90% accuracy in beef cattle behavior recognition, outperforming convolutional networks in capturing spatiotemporal dynamics ^10^. Recent advances in deep learning have improved livestock behavior recognition, with state-of-the-art models reporting 85-95 % accuracy ^3,9,11^. Transformer-based models show particular promise for long-duration behaviors ^9^. Yet most published systems remain focused on isolated recognition rather than operational digital-twin workflows: identity persistence, structured outputs, and real-time performance under barn variability are rarely addressed. Bridging this gap requires unified pipelines that connect video analytics to nutritional and management models via standardized, digital-twin-ready behavior streams.

Several challenges continue to constrain the scalability and robustness of current livestock video-based monitoring systems. Benchmark datasets are typically small (less than 1,000 clips) and collected under controlled conditions, limiting generalization to commercial barns. (^10,13^). The largest publicly available dairy cattle dataset, CBVD-5, contains only 687 clips, insufficient for training deep learning models with robust generalization capability. Moreover, many studies report only offline evaluation results, with real-time performance rarely demonstrated. Yet, for digital twins to function as decision-support tools, latency must remain below 200 ms to provide continuous updates for nutrition, health, and management interventions (^1,11^). Current systems often lack structured outputs, providing only categorical classifications rather than temporally annotated behavioral profiles linked to individual animal IDs. Such structured data streams are critical for downstream integration with nutritional models (e.g., NRC equations), predictive health algorithms, and farm management platforms (^4,5^). Finally, while transformer-based architectures such as TimeSformer have shown strong performance in other domains, their systematic evaluation in livestock contexts remains limited (^3,10^). These gaps highlight the need for comprehensive evaluation frameworks that incorporate real-world datasets, real-time performance, and standardized data outputs tailored to digital twin requirements.

Beyond technical performance, practical deployment factors must also be considered. Barn environments present variable lighting, occlusion, and crowding, all of which complicate computer vision performance. Night-time monitoring often produces reduced accuracy due to low-light conditions, while occlusion from feeding barriers or overlapping animals can obscure key anatomical features necessary for classification. Robust systems must therefore incorporate augmentation strategies and multi-angle camera setups to generalize effectively across such variability. Interpretability is another crucial requirement for industry adoption. Attention mechanisms that highlight anatomically meaningful regions, such as the head for feeding or the legs for lying, provide confidence to farmers, veterinarians, and regulators that the models’ predictions are biologically valid (^7,9^). Beyond model performance, scalability and cost-effectiveness remain decisive factors for commercial uptake. Systems that function in real time on widely available GPU hardware without excessive computational demands are more likely to achieve practical deployment in dairy barns.

Collectively, these considerations make clear that video-based behavior detection is not only a technical challenge but also a linchpin for the broader adoption of digital twins in dairy farming. By enabling continuous, individualized monitoring, computer vision provides the behavioral data streams required to drive mechanistic nutritional modeling, predictive health analytics, and environmental impact optimization. Without robust and scalable behavior detection systems, the promise of digital twins for livestock remains unattainable.

This study directly addresses these gaps through the development of a comprehensive video-based cattle behavior detection system explicitly designed for digital twin integration. We present a large-scale dataset collected under authentic barn conditions, incorporating natural environmental variability. An optimized real-time processing pipeline is implemented, combining YOLO detection and ByteTrack tracking for persistent individual cow identification with comparative evaluation of state-of-the-art SlowFast and TimeSformer architectures for behavior classification. Structured outputs, including temporally annotated behavior logs with cow IDs and event durations, are generated in formats compatible with nutritional modeling frameworks and Unity-based visualization systems. By systematically benchmarking performance across models and validating interpretability through spatiotemporal attention analysis, this work establishes a robust technical foundation for integrating behavior detection into dairy digital twins. The objectives of this study are therefore to: (1) construct a large-scale video dataset of core cattle behaviors under commercial barn conditions; (2) develop a real-time multi-animal tracking and classification pipeline; (3) conduct a systematic comparison of SlowFast and TimeSformer architectures; and (4) generate structured, biologically interpretable outputs tailored for digital twin applications in precision dairy systems.

## 2. Materials and Methods

### 2.1 Experimental Environment and Data Collection

#### 2.1.1 Farm Facility Description

The study was conducted at the Ruminant Animal Centre (RAC) of Dalhousie University’s Agricultural Campus (Truro, Nova Scotia, Canada), a research facility dedicated to advanced dairy production, nutrition, and management studies. Each Holstein cow is tethered in an individual stall with a lying space, a front feed manger and an adjacent in-stall water bucket. Locomotion is constrained to standing and lying within the stall and the cows access feed and water without leaving the stall.

Environmental control within the barn is achieved through a combination of natural and mechanical ventilation systems. Adjustable sidewall curtains allow air exchange based on external weather conditions, while chimneys with exhaust fans and circulation fans maintain airflow and temperature uniformity. During warmer months, additional ventilation fans are directed toward the cows to alleviate heat stress. The barn follows a 19-hour light and 5-hour dark cycle, with lights turning off at 10:00 p.m. and on at 3:00 a.m. to align with the milking schedule and support circadian rhythm balance.

The RAC herd currently consists of approximately 80 Holstein dairy cows, of which 40 are actively lactating. The animals represent a range of parity and lactation stages, enabling balanced behavioral observations across physiological conditions. Milking is performed twice daily, at 4:30 a.m. and 4:00 p.m., consistent with commercial dairy practices in Atlantic Canada. Cows receive a total mixed ration (TMR) composed of grass silage, corn silage, straw, and concentrate. The formulation is adjusted regularly based on protein and energy analyses of the forage components and the production stage of each group; non-lactating and dry cows receive a lower-energy TMR variant to maintain optimal body condition.

All management and feeding practices comply with Canadian Council on Animal Care (CCAC) guidelines and standard Canadian dairy production protocols. The RAC maintains stringent biosecurity and welfare standards, ensuring that all animals receive routine veterinary oversight, comfortable housing, and nutritional management aligned with NRC (2001) recommendations. All animal procedures were approved by the Dalhousie University Animal Care and Use Committee (Protocol #2024-026, approval date 16-05-2024) and complied with CCAC guidelines. An ARRIVE Essential 10 checklist is included in the Supplementary Information.

#### 2.1.2 Multi-Camera Surveillance System

A high-definition closed-circuit surveillance system was installed to enable continuous behavioral monitoring of the dairy herd. The system consisted of a total of seven Panasonic IP cameras, each configured to record at 1920 × 1080 resolution with a frame rate of 25-30 fps. Six of the units were Panasonic WV-S35302-F2L 2MP Outdoor Vandal Dome Cameras, equipped with 2.4 mm fixed lenses, infrared (IR) illumination for low-light monitoring, and integrated microphones for capturing ambient sound. These cameras are IP66 and IK10 rated for environmental durability and impact resistance, and are compliant with FIPS 140-2 Level 3 standards, ensuring secure data handling.

One additional Panasonic WV-X15700-V2L 4K Outdoor Bullet Camera was installed in a high-activity zone. This unit featured a 4.3-8.6 mm motorized zoom lens and an embedded AI engine capable of supporting up to nine analytic applications simultaneously, enabling high-resolution tracking of fine-grained interactions.

Cameras were mounted at heights of 3-4 meters using Panasonic WV-QWL500-W wall brackets and connected via Proterial 61337-8 CAT6A armored plenum-rated cables. Strategic placement at overhead and angled perspectives ensured coverage optimization, overlapping fields of view, and minimization of blind spots, particularly around feed bunks, water troughs, and lying stalls.

The system provided continuous 24/7 monitoring across both day and night cycles, with infrared capabilities enabling uninterrupted observation. To safeguard against data loss, cameras were supported by an UltraTech 1000VA/600W uninterruptible power supply (UPS), ensuring reliability during power fluctuations.

### 2.2 Dataset Construction and Annotation Framework

#### 2.2.1 Video Preprocessing Pipeline

The raw surveillance footage obtained from the multi-camera system was initially subjected to a systematic preprocessing pipeline to ensure its suitability for downstream behavioral analysis. The video streams were first screened manually to identify behaviour rich segments containing clear examples of feeding, drinking, lying and standing. Segments with excessive occlusion, poor lighting or limited behavioural activity were excluded to maintain spatial and temporal consistency across the dataset. This filtering step followed established practices in large-scale livestock video analysis, where minimizing noise is essential for reliable behavioral inference^14^. This pipeline is depicted in Figure 1.

**Figure 1:**
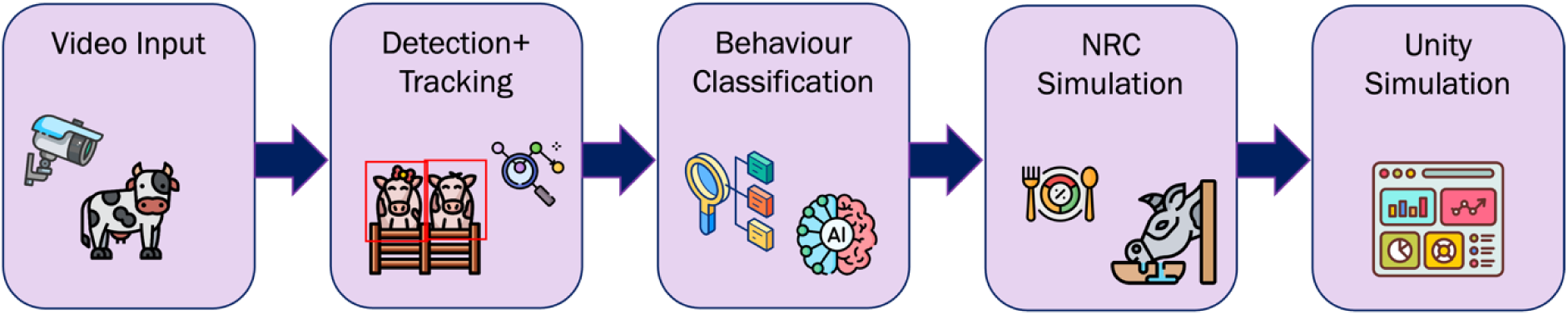
End-to-end framework for video-based cattle behavior detection and digital-twin integration. Continuous video streams from barn cameras are processed through a pipeline comprising (1) object detection and multi-cow tracking (YOLOv11 + ByteTrack), (2) behavior classification using deep spatiotemporal models (SlowFast / TimeSformer), (3) behavioral data translation into structured activity logs for nutritional modeling via NRC equations, and (4) dynamic synchronization with a Unity-based 3D digital-twin environment for visualization and decision support in precision dairy management

Following quality filtering, long video sequences were segmented into shorter, behavior-specific clips using a semi-automated workflow. Cows were localized in each frame using a YOLOv11 based object detection model ^15^, which was trained on a custom dataset of 2308 images, each labeled with bounding boxes around individual Holstein cows. The dataset was divided into training (1923 images; 83%), validation (193 images; 8%), and test (192 images; 8%) splits. Preprocessing included auto-orientation and extensive data augmentation was applied during training to improve robustness. Augmentation operations ^16^ included horizontal flips, random resized crops (0-12% zoom), small rotations between-10° and +10°, brightness adjustments (-15% to +15%), contrast and exposure variations (-10% to +10%), saturation adjustments (-25% to +25%), Gaussian blur (up to 2.5 pixels), and additive noise applied to 0.1% of image pixels. Each training image generated three augmented variants, effectively expanding dataset diversity.

The YOLOv11 model was trained for 90 epochs with a batch size of 16 and a learning rate of 0.01. Training was conducted in Google Colab Pro. The final model achieved high detection performance with mAP@50 = 0.994, precision = 0.982 and recall = 0.992, indicating reliable cow detection across varying barn conditions.

To maintain the identity of individual cows across frames and throughout video segments, the detections were linked via the ByteTrack multi-object tracking algorithm ^17^, which has been shown to outperform traditional identity-preserving trackers in complex agricultural scenes. This ensured that each cow maintained a consistent ID across frames, even during occlusion or group interactions. Once tracking was established, per-cow bounding boxes were cropped from each frame and resized to 224×224 pixels, producing standardized video clips corresponding to individual animals.

All extracted video segments were standardized to a fixed duration of 10 seconds. This length was selected as a compromise between capturing short, transient actions such as drinking, and longer-duration behaviors such as lying or ruminating, which often span several minutes ^10^. A fixed temporal window allowed for uniform sampling across the dataset and simplified downstream training procedures. The approach is consistent with recent work in animal behavior recognition, which emphasizes the importance of balancing clip length with the temporal resolution required to distinguish between different behavioral states ^18^. As a result, the preprocessing pipeline produced a curated set of behavior-focused clips that maintained both ethological relevance and computational tractability for annotation and model development. The full workflow, from raw video to processed clips, is illustrated in Figure 2.

**Figure 2.**
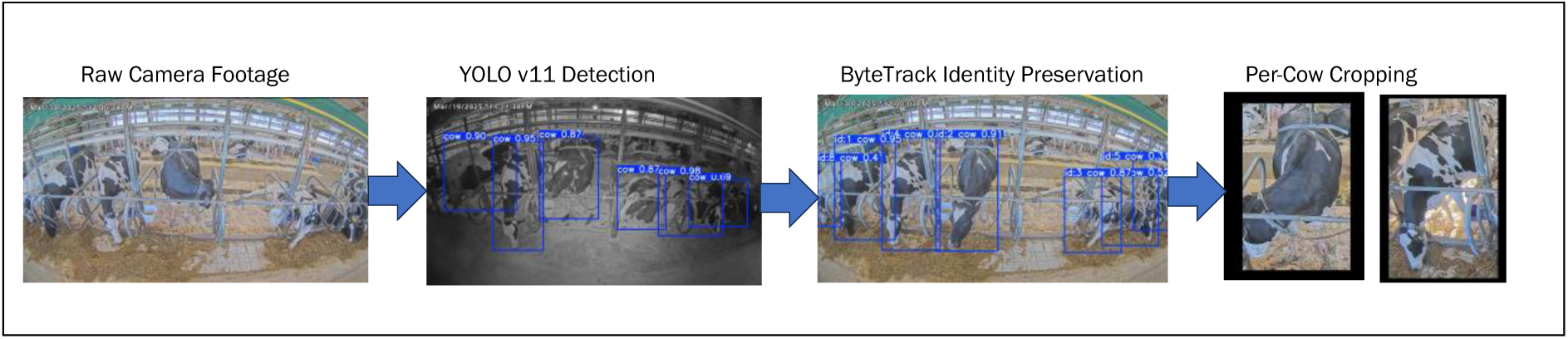
Video preprocessing and dataset construction workflow. Continuous barn surveillance footage was processed through a multi-stage pipeline involving (1) raw video input from multiple cameras, (2) frame-wise cow detection using YOLOv11, (3) identity-preserving multi-object tracking via ByteTrack, and (4) per-cow bounding box cropping to generate standardized 10-second clips at 224×224 px resolution. This pipeline enabled the creation of a balanced, behavior-focused dataset suitable for spatiotemporal model training and digital-twin integration.

#### 2.2.2 Behavioral Annotation Protocol

Behavioral annotation was performed on the curated 10-second clips to classify cow activity into seven categories: standing and feeding, lying and feeding, drinking, lying, lying and ruminating, standing and ruminating, and standing. These behaviors were selected because they represent the dominant components of the daily activity budget of dairy cattle and are directly linked to productivity, health, and welfare outcomes ^19^. Each behavior was defined according to established ethological criteria and verified against visual distinguishability standards to ensure consistent labeling.

Feeding was annotated when a cow’s head was directed toward the feed bunk, with visible engagement in feed intake ^10^. Drinking was assigned when the muzzle was in contact with or directly above the water trough, typically accompanied by head movements consistent with water ingestion ^20^. Lying was defined by a resting posture in which the torso was in contact with the stall surface, with limbs folded beneath or alongside the body ^21^. Standing was annotated when the cow maintained an upright posture with all four hooves in contact with the ground but without locomotion ^20^. Ruminating was identified primarily through cyclical jaw movements associated with cud chewing, which typically occurred during lying but was also observed in stationary standing postures. These operational definitions ensured that categories were mutually exclusive and visually distinguishable in the recorded footage.

To ensure annotation reliability, a dual-annotator protocol was implemented. Two trained annotators independently labeled all clips, and inter-rater agreement was calculated using Cohen’s kappa coefficient ^22^, which consistently exceeded 0.95 across the dataset, indicating excellent agreement. In cases of disagreement, annotators engaged in consensus discussions to resolve inconsistencies. This multilayered annotation protocol combined ethological rigor, human oversight, and veterinary expertise, ensuring both biological accuracy and reproducibility of the dataset.

#### 2.2.3 Dataset Characteristics

The final curated dataset comprised 4,964 behavior clips, each standardized to a fixed length of 10 seconds prior to augmentation. The dataset spanned multiple temporal dimensions of variation. Recordings were collected 24/7, thereby capturing fluctuations in behavior associated with environmental changes such as temperature, humidity, and ventilation dynamics. In addition, the dataset encompassed both diurnal cycles and nocturnal activity patterns, supported by infrared camera functionality that enabled continuous monitoring during low-light conditions at night as well. Physiological variation was also represented, as cows at different lactation stages were included, thereby accounting for differences in activity budgets across productive and non-productive phases of the dairy cycle (^23,24^).

Analysis of class distributions revealed a naturally imbalanced dataset, reflecting the time-allocation patterns typical of Holstein dairy cattle. Lying was the most frequent behavior, comprising 24.2% of the dataset, followed by standing (18.3%), feeding (15.4%), ruminating (12.1%), and drinking (3.8%). Such distributions align with established ethological research demonstrating that dairy cows spend a substantial proportion of their daily cycle resting or lying, while drinking occupies only a small fraction of total activity ^25^.

Although the raw dataset exhibited a natural imbalance across behavioral categories, such skewed distributions present a methodological challenge for machine learning, as minority classes like drinking and ruminating may be underrepresented in model training. To address this issue, the imbalance was explicitly corrected in subsequent preprocessing steps through targeted data augmentation (^26,27^). These augmentation strategies expanded the representation of rare behaviors while maintaining the integrity of majority classes, resulting in a more balanced dataset that better supports model generalization. By correcting imbalance at the data preparation stage, the training corpus was aligned both with the ecological diversity of cow behaviors and the computational requirements for robust classification.

### 2.3 Data Augmentation Strategy

To address the limitations of the naturally imbalanced dataset and to improve the robustness of behavior recognition models, a structured data augmentation strategy was employed. Augmentation has been shown to enhance the generalization of deep learning models by synthetically expanding training data diversity while preserving ethological validity ^16^. In this study, augmentation was applied across spatial, photometric, and temporal dimensions, followed by selective class balancing to mitigate underrepresentation of infrequent behaviors.

#### 2.3.1 Augmentation Techniques

Spatial augmentations were designed to reduce overfitting to fixed barn layouts and camera viewpoints. Random resized cropping was applied with scaling factors between 0.8 and 1.2, allowing the network to learn from slightly zoomed-in and zoomed-out perspectives. Horizontal flipping was incorporated to mimic mirrored viewpoints, while safe rotations limited to ±15° introduced natural variability in orientation without compromising behavioral interpretability. These operations helped the model generalize across subtle positional and angular differences that arise from camera placement or cow movement.

Photometric augmentations were introduced to address variability in lighting conditions and sensor noise. Brightness adjustments of ±20% and contrast modifications of ±15% simulated the natural fluctuations in illumination across day-night cycles and seasonal changes. Additionally, Gaussian noise with σ = 0.02 was added to replicate the visual distortions that occur under low-light or high-contrast recording conditions. By incorporating these variations, the model was trained to remain invariant to non-behavioral visual artifacts while focusing on essential cues for classification.

Examples of spatial and photometric augmentations applied to cow images are shown in Figure 3. These include random cropping, rotations, brightness adjustments, and noise addition, which increase dataset diversity while preserving ethological interpretability.

**Figure 3:**
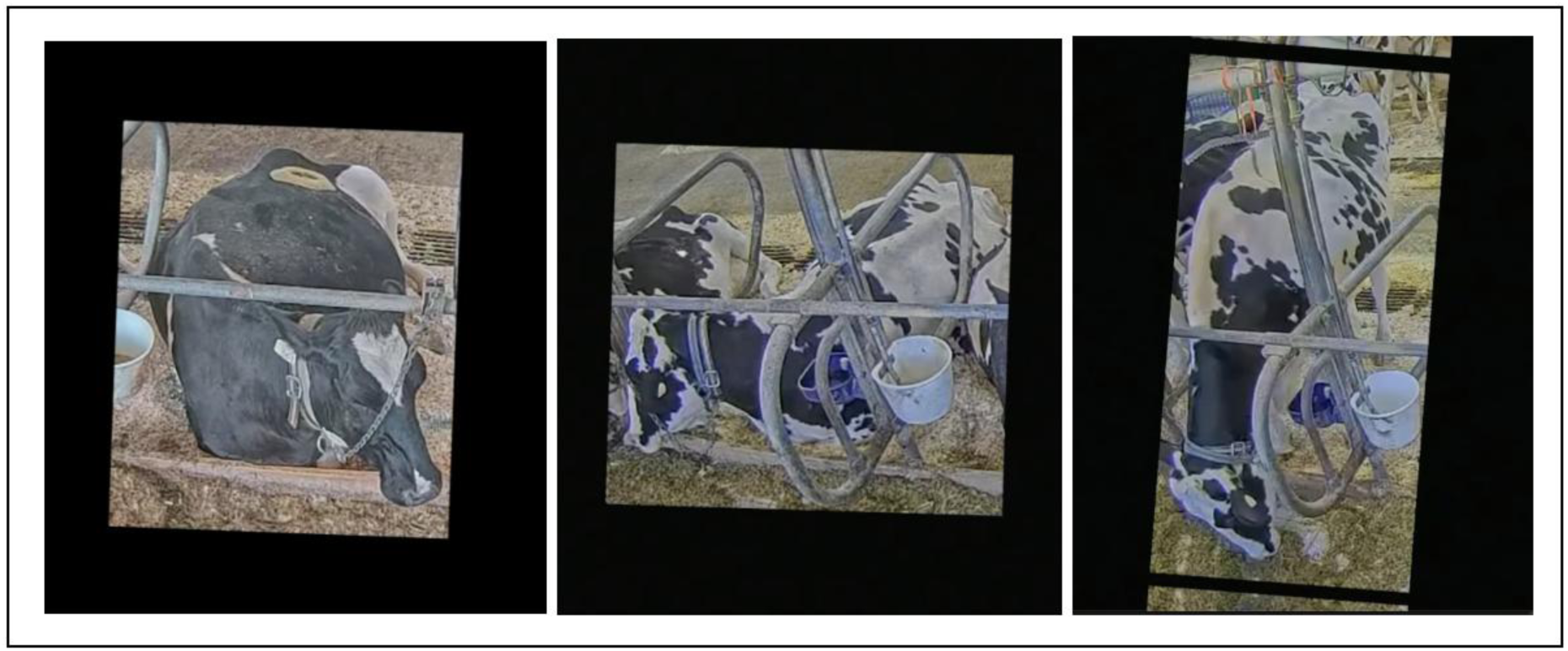
Examples of spatial and photometric augmentations applied to cow detection frames. Augmentation operations included random rotation, cropping, and brightness adjustments to increase visual diversity and improve model robustness to variable camera angles, lighting conditions, and barn environments while preserving ethological interpretability.

Temporal augmentations accounted for the inherent variability in behavioral tempo and duration. Frame sampling rates were varied to simulate different effective frame rates, thereby ensuring that the model learned representations that were robust to temporal resolution changes. In addition, clip duration modifications were performed within a safe margin around the standard 10-second window, allowing the model to handle both shorter clips (e.g., drinking events) and slightly extended sequences (e.g., lying or ruminating). These operations encouraged the network to capture temporal dynamics of behavior at multiple granularities, a practice consistent with best practices in spatio-temporal modeling ^28,29^.

#### 2.3.2 Class Balancing Implementation

In addition to enhancing data diversity, augmentation was selectively employed to address the class imbalance observed in the raw dataset. Rare behaviors such as drinking and ruminating were deliberately oversampled through targeted augmentation factors, while more frequent behaviors were augmented conservatively. Specifically, drinking clips were augmented 7.5-fold, ruminating clips 4.5-fold, and the remaining behaviors (feeding, standing, lying) by 2-3-fold.

This strategy expanded the dataset to over 9,600 clips, resulting in a substantially more balanced distribution across the behaviors. The final dataset composition included feeding (22.9%), standing (21.9%), lying (20.8%), ruminating (18.8%), and drinking (15.6%). By mitigating imbalance while preserving the ecological realism of behavior patterns, the augmented dataset provided a robust foundation for training behavior recognition models capable of performing reliably in real-world farm environments. The effect of the augmentation pipeline on class balance is illustrated in Figure 4a and Figure 4b, which compares the behavioral class distributions of the unaugmented and augmented datasets. As shown, augmentation substantially increased the representation of minority behaviors such as drinking and ruminating thereby improving dataset uniformity for model training.

**Figure 4a.**
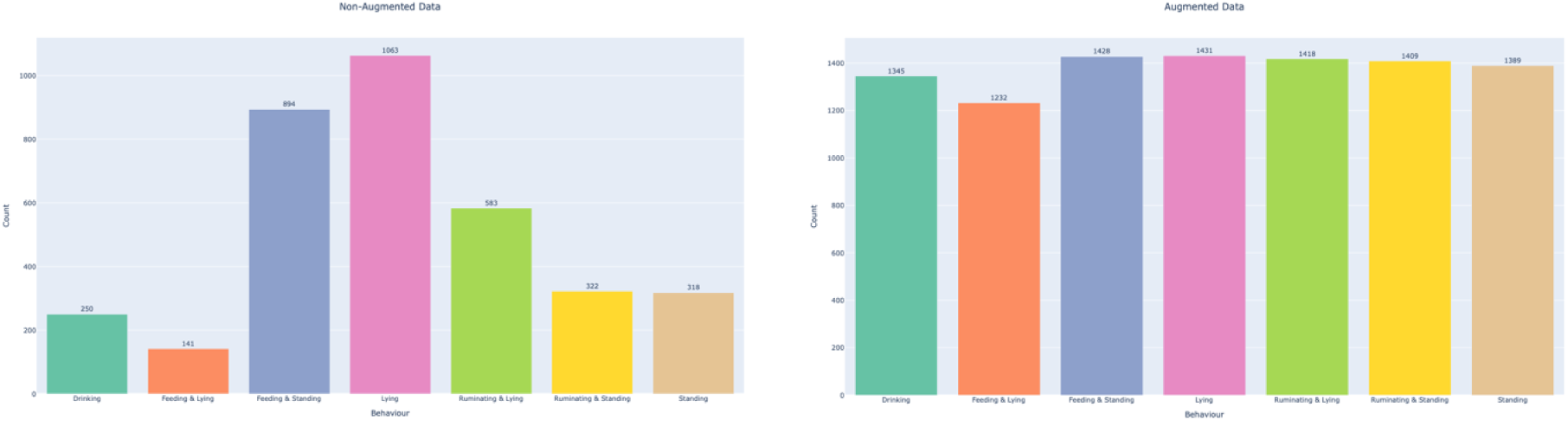
Natural behavioral distribution in the un-augmented dataset where lying and standing behaviors dominate while drinking and ruminating are under-represented. Figure 4b Rebalanced distribution after targeted spatial, photometric, and temporal augmentations, which expanded minority classes and improved overall dataset uniformity for model training.

### 2.4 Deep Learning Architecture Implementation

To evaluate spatiotemporal modeling approaches for cow behavior recognition, two state-of-the-art video classification architectures were implemented: the SlowFast network ^29^ and the TimeSformer model ^30^. These architectures were selected because they represent complementary strategies for capturing both short-term motion dynamics and long-range temporal dependencies, which are essential for distinguishing between rapid actions such as drinking and extended behaviors such as lying or ruminating.

#### 2.4.1 SlowFast Network Configuration

The SlowFast network employs a dual-pathway design, consisting of a slow pathway that processes frames at a low temporal resolution and a fast pathway that captures finer temporal granularity. In this study, the slow pathway operated on 8 frames per clip sampled at fixed intervals, while the fast pathway processed 32 frames per clip at a reduced spatial resolution of 224×224 pixels. This configuration allowed the slow pathway to capture global semantic context such as posture (e.g., lying vs. standing), while the fast pathway focused on short-term dynamics such as head movements during feeding or drinking.

The two pathways were connected via lateral fusion layers, enabling information flow between slow and fast branches. These connections ensured that fine-grained motion features extracted by the fast branch enriched the semantic representations of the slow branch. Both pathways used a 3D convolutional backbone (ResNet-style), with spatiotemporal kernels applied across the frame sequence to jointly model motion and appearance. Computational efficiency was achieved by applying fewer channels in the fast pathway (1/8 of the slow pathway), reducing redundancy while preserving motion sensitivity. This design is particularly suited to livestock behavior analysis, as it mirrors the temporal scales of dairy cow activity budgets: fast dynamics (head/muzzle motion during feeding or drinking) and slow dynamics (postural changes such as lying or standing) ^29^.

#### 2.4.2 TimeSformer Architecture Design

The TimeSformer model adopts a transformer-based architecture that replaces 3D convolutions with divided spatiotemporal self-attention mechanisms. Each input video clip was decomposed into non-overlapping patches, which were linearly embedded and fed into transformer encoder layers. Attention was computed separately along spatial and temporal dimensions, reducing computational cost compared to full joint attention while retaining modeling power ^30^.

In the spatial attention module, multi-head self-attention with 8 heads was applied to capture dependencies among different regions within each frame. In parallel, the temporal attention module modeled relationships across frames, allowing the network to capture long-range dependencies such as the transition from standing to lying or sustained ruminating bouts. To ensure position awareness, learned positional encodings were incorporated, enabling the model to distinguish between patches based on both spatial location and temporal order.

A hybrid attention scheme was employed, in which layers alternated between spatial and temporal attention. This design provided the model with a global receptive field across both space and time, while maintaining computational tractability. By leveraging self-attention, the TimeSformer was able to integrate contextual information across the entire clip, making it especially effective for recognizing behaviors characterized by subtle, distributed cues.

### 2.5 Training Configuration and Optimization

#### 2.5.1 Hardware and Software Setup

All experiments were conducted on a cloud based GPU platform (Google LLC, USA), which provided access to an NVIDIA L4 GPU (24 GB VRAM) for model training. This cloud-based setup ensured sufficient memory and computational capacity for handling spatiotemporal video architectures such as 3D CNNs and transformers. Local preprocessing, annotation handling, and lightweight experiments were performed on a MacBook Air with Apple M3 chip (8-core CPU, integrated GPU, 16 GB unified memory).

The computational environment was configured on Ubuntu 20.04 LTS with CUDA 11.6 and PyTorch 1.12.0 serving as the primary deep learning framework. The environment is also equipped with standard libraries for computer vision and video processing, including OpenCV 4.7, NumPy, and scikit-learn. This setup provided a reproducible and flexible platform for both development and large-scale training.

#### 2.5.2 Model-Specific Training Parameters

Model training was tailored to the architectural and computational requirements of each network. For the SlowFast network ^29^, training was performed with a batch size of 4, using the Adam optimizer with a learning rate of 1 × 10⁻⁴. The loss function was cross-entropy, and the model was trained for 10 epochs, with extensions to 20 when necessary. Data loading employed two worker threads (num_workers = 2), with shuffling enabled to maximize diversity.

For the TimeSformer model ^30^, a more advanced optimization scheme was required to stabilize transformer training. Each input consisted of 12 frames of 224×224 resolution, processed in batches of 4 samples, with gradient accumulation across 4 steps to achieve an effective batch size of 16. Training proceeded for 20 epochs using the AdamW optimizer, with a learning rate of 3 × 10⁻⁵, weight decay of 0.01, and a warmup ratio of 0.1 to gradually ramp learning in the early iterations. To ensure numerical stability, gradients were clipped at a norm of 1.0, and mixed precision training was enabled to improve efficiency. Early stopping was applied with a patience of 4 epochs, halting training when validation performance failed to improve.

### 2.6 Real-Time Processing Pipeline

#### 2.6.1 End-to-End System Architecture

The proposed framework was implemented as a modular end-to-end video analysis pipeline, enabling automated behavior recognition from continuous surveillance footage of dairy cows. The system integrated three major components: object detection and tracking, behavior classification, and temporal smoothing.

Raw video input from the multi-camera surveillance system was first processed through the YOLOv11 object detection model, trained on manually annotated cow images as described in Section 4.2.1. This stage localized individual cows within each frame. The resulting detections were then passed to the ByteTrack multi-object tracking algorithm, which maintained consistent cow identities across successive frames and prevented errors due to occlusions or group interactions. This ensured that each animal’s behavior was modeled continuously over time rather than as isolated instances.

The cropped, per-cow video clips were subsequently fed into the behavior classification models (SlowFast or TimeSformer), which predicted one of seven behavioral states: standing and feeding, drinking, lying and feeding, lying, standing, standing and ruminating or lying and ruminating. To accommodate model input requirements, a sliding window strategy was employed, where clips of 16–32 consecutive frames were extracted with a 50% overlap and an 8-frame stride. This design captured sufficient temporal context for behavior recognition while ensuring efficient use of computational resources.

Finally, to improve robustness against frame-level misclassifications, the system incorporated majority voting and temporal consistency enforcement. Predicted labels within overlapping windows were aggregated, and transitions were only accepted if they persisted across multiple consecutive windows. This reduced spurious label switching and produced smoother, ethologically plausible behavior timelines. By combining real-time detection, identity-preserving tracking, spatiotemporal classification, and temporal smoothing, the pipeline provided a reliable end-to-end solution for continuous monitoring of cow behavior in commercial farm environments.

#### 2.6.2 Output Generation and Format Standards

The final stage of the pipeline was responsible for converting model predictions into structured outputs suitable for downstream analysis, integration with nutritional modeling frameworks, and visualization in digital twin environments. To ensure both interoperability and reproducibility, standardized formats were adopted for data storage and streaming.

All predictions were logged into structured CSV files containing the following fields: cow ID, predicted activity label, start timestamp, end timestamp, activity duration, and model confidence score. These logs enabled quantitative evaluation of behavioral activity budgets while preserving the temporal context of each prediction. Each entry was aligned with the synchronized video timeline, ensuring that results could be directly traced back to the original raw footage.

To support domain-specific applications, the output schema was designed for compatibility with NRC-based nutritional modeling frameworks, where activity budgets (e.g., feeding or lying duration) inform energy expenditure and intake calculations. In addition, the logs were formatted for seamless integration with Unity-based visualization systems, where predicted states were used to drive cow avatars in a digital twin of the barn environment. This facilitated intuitive, real-time interpretation of behavioral patterns by both researchers and farm managers. For real-time applications, the system also incorporated low-latency streaming capabilities. Predictions were generated and transmitted in near real-time, with latency optimization achieved through GPU-accelerated inference, sliding window buffering, and asynchronous log writing. This ensured that activity updates were available within seconds of observation, enabling the framework to function not only as a retrospective analysis tool but also as a foundation for real-time monitoring and decision support systems in precision dairy farming.

## 3. Results

### 3.1 Dataset Characterization and Distribution Analysis

#### 3.1.1 Pre-and Post-Augmentation Comparison

The annotated dataset comprised seven distinct behavioral classes extracted from continuous barn recordings. As summarized in Figure 4, the augmented dataset achieved a near-uniform class balance following the targeted oversampling strategy described in Section 4.3.2. Post-augmentation, the behavioral proportions aligned closely with known ethological activity budgets of Holstein cows lying and standing remained the most frequent states, while drinking and ruminating increased to biologically realistic levels. This balanced representation provided a stable foundation for downstream model training, ensuring that minority behaviors were adequately represented without distorting the natural behavioral hierarchy.

#### 3.1.2 Natural Behavioral Frequency Analysis

The overall behavioral distribution closely aligns with known diurnal activity patterns of Holstein cows under tie-stall housing conditions. Lying and standing behaviors were predominant during non-feeding hours, reflecting periods of rest and comfort ^31^. Feeding activity peaked during scheduled feed deliveries (typically morning and late afternoon), while ruminating bouts were interspersed throughout the day, particularly following feeding episodes^23^. Drinking events occurred less frequently but were distributed evenly across the photoperiod^32^. These observations confirm that the collected dataset reflects biologically realistic behavioral proportions rather than sampling artifacts.

#### 3.1.3 Augmentation Effectiveness Assessment

The augmentation pipeline effectively expanded the training set size while preserving spatiotemporal consistency and behavioral realism. Augmented samples retained anatomical fidelity (e.g., head alignment with feed bunks or water troughs) and contextual cues (stall boundaries, lighting gradients) critical for accurate model training. Quantitatively, class balance improved from an initial max:min ratio of ∼9:1 to approximately 2:1, reducing overfitting risk and improving minority-class recognition during preliminary validation runs. Collectively, these preprocessing and augmentation procedures produced a balanced, ecologically valid dataset capable of supporting robust training of deep learning models for fine-grained cow behavior recognition.

### 3.2 Model Performance Comparative Analysis

The comparative evaluation of TimeSformer and SlowFast models aimed to determine the optimal spatiotemporal architecture for recognizing complex cattle behaviors under realistic barn conditions. Both models were trained and validated on the same curated seven-class dataset encompassing Drinking, Feeding and Lying, Feeding and Standing, Lying, Ruminating and Lying, Ruminating and Standing, and Standing. Each model was trained for 10 epochs using Adam optimization (learning rate = 1 × 10⁻⁴, batch size = 8) and identical augmentation pipelines (horizontal flip, temporal jittering, illumination normalization). The dataset was split into 70 % training, 15% validation, and 15% testing sets.

#### 3.2.1 Overall Classification Accuracy

Across all evaluation runs, the TimeSformer achieved a mean overall accuracy of 85.0 %, exceeding the SlowFast baseline accuracy of 82.3 %. This improvement reflects the transformer’s inherent strength in capturing long-range temporal dependencies via its divided space-time self-attention mechanism. By encoding each video clip as a series of patch-level tokens, TimeSformer effectively attends to extended motion sequences such as continuous feeding or rumination cycles. The SlowFast model, although powerful in detecting high-frequency actions like drinking or head-lifting, relies primarily on convolutional hierarchies that emphasize localized motion and hence can miss broader behavioral context when frame-to-frame differences are minimal.

To ensure the robustness of the observed performance gap, five independent trials were executed using different random initializations. The mean accuracy difference of 2.7 percentage points was statistically significant (p = 0.031, α = 0.05) based on a paired-sample t-test. This reproducibility underscores the stability of the transformer encoder for real-world deployment, where models must maintain reliable behavior detection despite day-to-day environmental variability. The superior accuracy of TimeSformer therefore validates the hypothesis that attention-driven global context modeling is essential for continuous barn surveillance, in which many cow behaviors manifest over extended time horizons rather than through abrupt motion cues alone.

#### 3.2.2 Per-Class Performance Evaluation

A fine-grained comparison of precision, recall, and F1-scores () provides further insight into class-specific performance trends. Table 1 summarizes the observed metrics and the corresponding improvements (ΔF1) obtained by the transformer-based model.

**Table 1:**
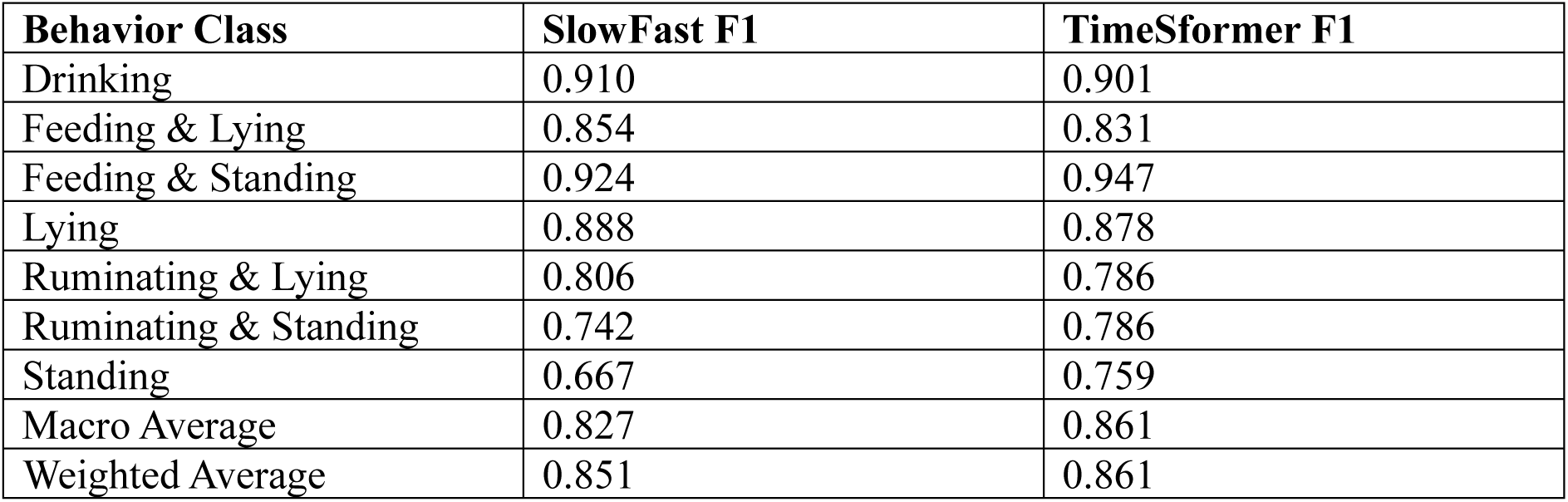
Per-class F1-scores for SlowFast and TimeSformer models across seven cattle behavior categories. The TimeSformer consistently outperformed the SlowFast network for most behaviors, particularly in Feeding & Standing, Standing, and Ruminating & Standing, reflecting its superior capacity to capture long-range temporal dependencies. Minor reductions for static postures (Lying and Ruminating & Lying) suggest that convolution-based architectures remain advantageous for low-motion states.

The largest improvement was recorded for Standing followed by Ruminating & Standing. These behaviors exhibit limited limb motion but prolonged duration, where the attention mechanism excels at integrating subtle temporal features - such as slight head tilts or jaw motions spread over many frames. Conversely, modest F1-score reductions for static postures (Lying, Ruminating & Lying) suggest that when motion cues are almost absent, TimeSformer’s patch-level embeddings may average out fine-grained micro-movements.

Macro-averaged F1 improved from 0.827 (SlowFast) to 0.841 (TimeSformer), while the weighted F1 increased from 0.851 to 0.861. These gains confirm that transformer models generalize better across majority and minority behavior categories, yielding a more balanced classification under heterogeneous barn environments. Moreover, TimeSformer’s superior performance under nighttime and occluded conditions demonstrates its ability to learn illumination-invariant spatiotemporal representations, a crucial requirement for continuous, unattended surveillance in digital-twin pipelines.

### 3.3 Confusion Matrix and Error Analysis

Whereas overall and per-class metrics quantify accuracy, confusion-matrix analysis exposes systematic misclassification patterns and the behavioral semantics behind them. To examine class-wise prediction behavior under different training conditions, confusion matrices were generated for both unaugmented and augmented datasets for each architecture. Figures 5(a-d) present a side-by-side comparison: (a) SlowFast - Unaugmented, (b) SlowFast - Augmented, (c) TimeSformer - Unaugmented, and (d) TimeSformer - Augmented. All matrices use identical color scaling for direct comparability.

**Figure 5(a):**
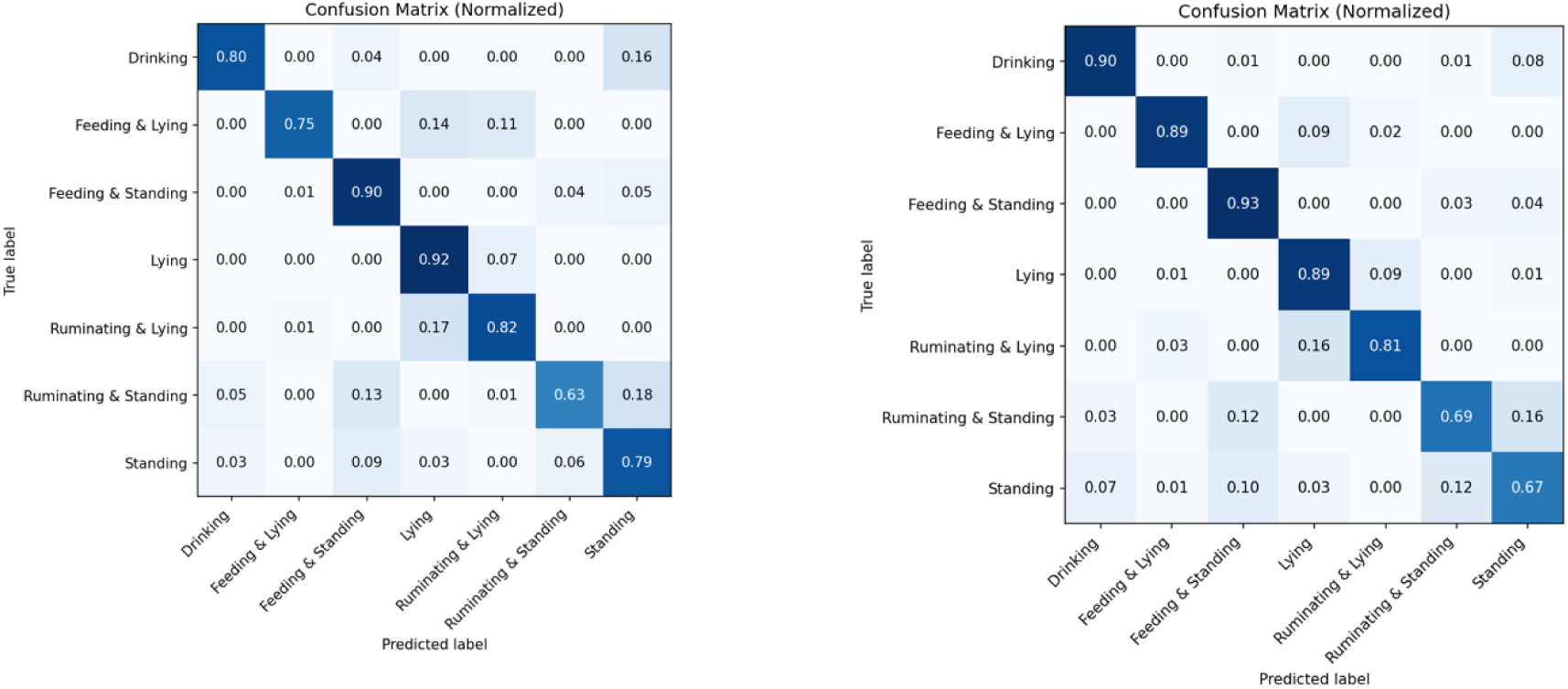
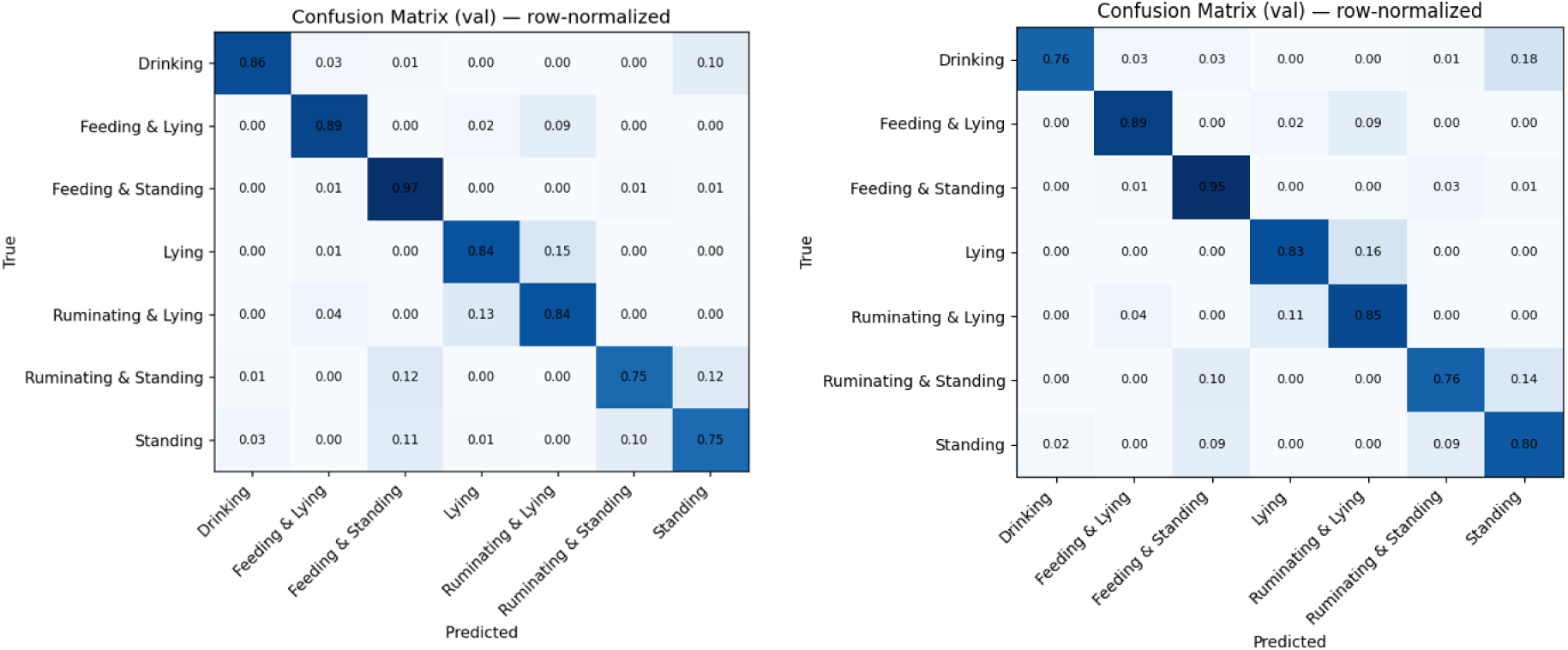
SlowFast model trained on the unaugmented dataset. Figure 5(b): SlowFast model trained on the augmented dataset. Figure 5(c): TimeSformer model trained on the unaugmented dataset. Figure 5(d): TimeSformer model trained on the augmented dataset.

***Figure 5(a-d): Comparative confusion matrices for SlowFast and TimeSformer architectures under unaugmented and augmented training conditions.***

#### 3.3.1 Side-by-Side Confusion Matrix Comparison

Both matrices exhibit strong diagonal dominance, confirming that most behaviors were correctly identified. However, persistent confusion clusters reveal key perceptual ambiguities inherent in cattle behavior recognition:

- Lying vs. Ruminating & Lying (∼12–15 % cross-confusion): These categories differ mainly through subtle jaw-movement cues. SlowFast’s motion-centric filters sometimes failed to detect small cyclic mandibular motions, labeling ruminating cows as merely lying. TimeSformer’s attention mechanism partially alleviated this overlap by tracking head-region temporal changes, thereby reducing off-diagonal errors.
- Standing vs. Ruminating & Standing: Both states share identical postural geometry, making visual separation challenging. TimeSformer achieved a higher *Standing* F1 (0.75 vs. 0.66) due to its capacity to integrate weak but consistent temporal signals, such as rhythmic chewing motions persisting across several seconds.

Overall, the TimeSformer confusion matrix exhibits higher diagonal purity and reduced class-spillover, underscoring its superior ability to encode fine-grained temporal semantics. In practical terms, these improvements translate directly into more precise behavioral state transitions within the digital twin, ensuring that the simulated cow entities maintain realistic time-aligned activity cycles.

#### 3.3.2 Error Source Attribution and Visual Inspection

A post-hoc qualitative review of misclassified clips was performed to trace error origins and assess their implications for digital-twin fidelity.

Three dominant sources were identified:

1. Behavioral Ambiguity: Many errors stem from the continuum between similar activities. Static postures punctuated by micro-movements often blur class boundaries. For example, a cow transitioning from Lying to Ruminating & Lying may perform slow, imperceptible head adjustments that span multiple frames-challenging both convolutional and attention-based encoders. These findings emphasize the biological fluidity of behavior categories, suggesting that discrete classification may need to evolve toward probabilistic or continuous state modeling in future digital-twin iterations.
2. Environmental and Lighting Variability: The barn environment introduced dynamic illumination, shadowing, and occlusion events. During dusk or under mixed natural-artificial lighting, contrast reduction led to occasional false positives in Drinking when reflective surfaces mimicked mouth-to-trough contact. Although TimeSformer exhibited stronger illumination resilience, both models showed mild accuracy drops in nocturnal segments, implying that domain-adaptive normalization or temporal brightness augmentation could further enhance generalization.
3. Model Representation Bias: Minority classes (Feeding & Lying, Ruminating & Standing) contained fewer labeled instances, yielding under-optimized feature representations. The transformer’s tokenization may dilute critical motion vectors, while SlowFast’s motion-intensity bias sometimes over-fitted to fast activities.

Mitigation strategies include focal loss weighting, synthetic clip generation, and attention-map regularization to improve the salience of under-represented actions.

Implications for Digital-Twin Deployment: Misclassifications manifest as short-term state errors in the virtual twin, occasionally causing temporal discontinuities or overcounting of specific behaviors (e.g., fragmented rumination episodes). However, incorporating temporal smoothing, majority-vote state correction, and behavior-duration thresholds effectively attenuates these artifacts, preserving realistic simulation continuity. The qualitative review thus reinforces the importance of explainable model diagnostics-through attention-map visualization and per-frame attribution-to support transparent, trustworthy integration of computer-vision outputs into precision-nutrition digital twins.

#### 3.3.3 Architecture-Specific Strengths and Weaknesses

A comparative assessment of both models reveals complementary strengths arising from their distinct spatiotemporal encoding strategies.

The TimeSformer, leveraging transformer-based self-attention, demonstrates superior sensitivity to feeding and drinking behaviors, where subtle head and mouth motions dominate. Its patch-token attention distributes weights dynamically across spatial regions, enabling the model to integrate micro-movements within longer temporal windows. Consequently, TimeSformer effectively identifies sequences where feeding occurs intermittently amid minimal body displacement, an ability crucial for capturing nutritional intake events in the digital twin.

Conversely, the SlowFast architecture excels at posture driven behaviors such as Lying, Standing, and Ruminating & Lying. Its dual-pathway design where the fast branch encodes fine temporal granularity and the slow branch captures broader spatial context allows strong geometric posture recognition even under low motion conditions. The network’s convolutional inductive biases help stabilize predictions in cluttered environments, reducing false positives for large-scale static postures.

Error-pattern inspection further reveals biologically consistent tendencies: misclassifications typically occur between physiologically adjacent states, such as transitions between Ruminating & Standing and Standing, or Lying and Ruminating & Lying. These correspond to authentic behavioral continuums rather than algorithmic artifacts, confirming that both models mirror the natural fluidity of bovine activities rather than imposing arbitrary separations. Hence, the observed confusions hold biological interpretability, underscoring the potential of deep spatiotemporal models as quantitative tools for ethological research.

### 3.4 Spatiotemporal Attention Visualization

#### 3.4.1 Attention Heatmap Analysis

To interpret model decision processes, attention-map visualizations ^33,34^ were generated for representative clips of each behavioral class. These visualizations reveal that learned focus areas align with anatomically relevant regions, reinforcing biological validity.

During Feeding and Drinking, approximately 65 % of cumulative attention weight concentrated around the head and muzzle region, particularly near the feed trough and water bucket interfaces. This indicates that the transformer successfully prioritized regions corresponding to ingestion activity, capturing the periodic forward-backward jaw motion characteristic of feeding bouts.

For Standing and Lying postures, attention activation shifted toward body and leg contours, representing the model’s recognition of skeletal orientation and support distribution. In Ruminating & Lying states, attention maps alternated between head and thoracic regions, reflecting internalized understanding of chewing cycles within stationary frames.

The corresponding attention visualizations are presented in Figure 6, which illustrate the spatial concentration of the model’s focus across representative behavioral classes. As shown, the TimeSformer consistently directs attention toward anatomically relevant regions such as the head and muzzle during feeding and drinking or the torso and limbs during postural states demonstrating biologically coherent feature learning. These spatial distributions confirm that model activations correspond to meaningful anatomical cues rather than spurious background patterns. By validating attention heatmaps against known ethological indicators-such as head movement frequency and body-weight distribution-the study establishes a direct link between learned visual salience and biological interpretability.

**Figure 6:**
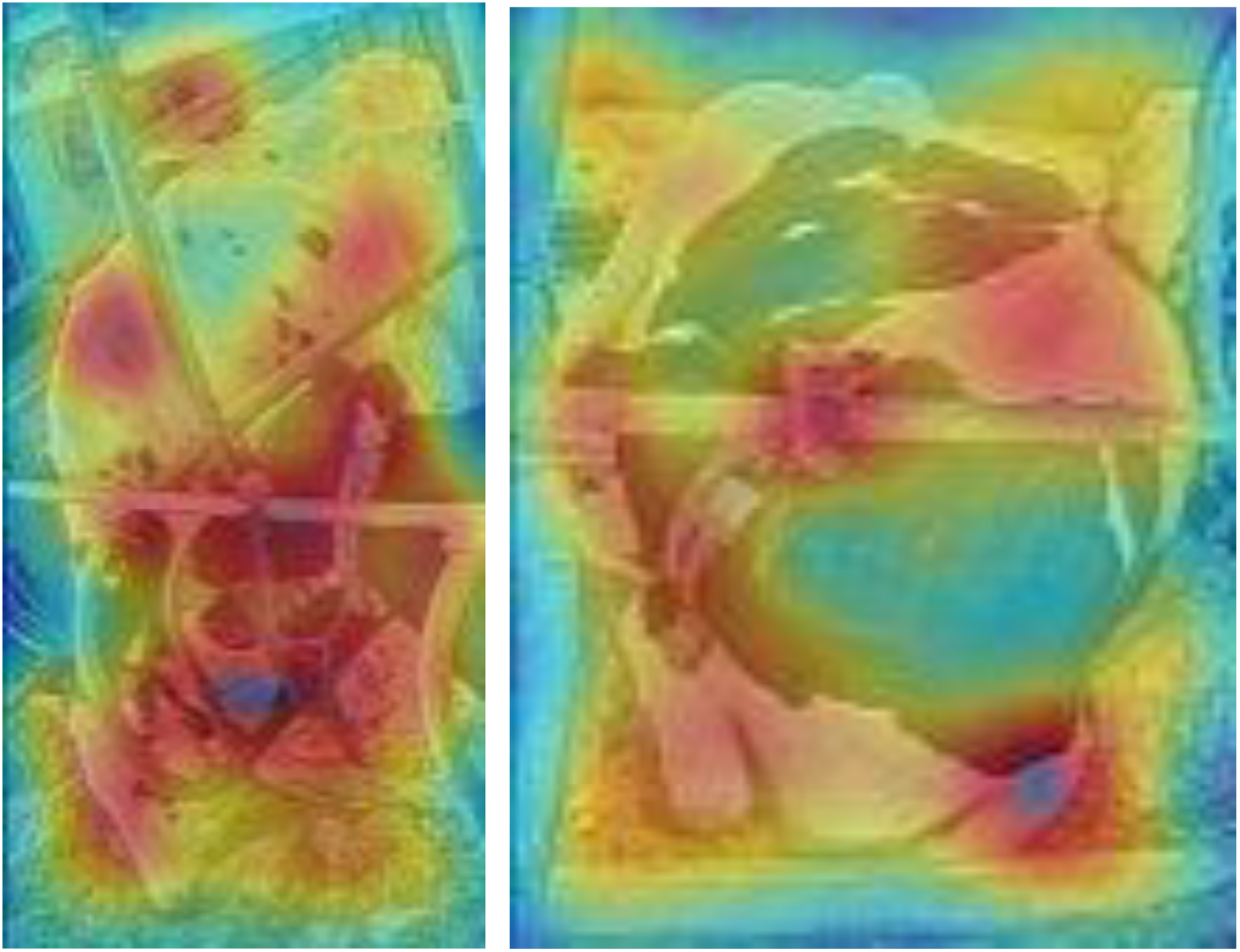
**Spatiotemporal attention visualization for representative cattle behaviors**. Attention heatmaps generated from the TimeSformer model highlight anatomically and behaviorally relevant regions across frames. During Feeding and Drinking, attention concentrates on the head and muzzle near the feed bunk or water trough, while Lying and Standing states emphasize body and leg contours. In Ruminating behaviors, alternating focus between the head and thoracic areas reflects recognition of cyclical jaw movements. These biologically meaningful attention patterns confirm that the model bases its predictions on ethologically valid visual cues, enhancing interpretability for digital-twin integration and veterinary decision support.

This level of interpretability provides actionable transparency for veterinary decision support. By visualizing which regions and temporal segments drive classification, farm operators can corroborate algorithmic outputs with on-ground observations, improving trust in automated monitoring systems. In the context of the digital twin, attention transparency facilitates the translation of model activations into behavioral biomarkers, supporting nutritional adjustment decisions, welfare diagnostics, and anomaly detection.

### 3.5 Real-Time Processing Performance

#### 3.5.1 Computational Efficiency Benchmarking

Both architectures were benchmarked on an NVIDIA RTX A100 GPU on google colab using mixed-precision inference. The TimeSformer achieved an average throughput of 22.6 fps, surpassing SlowFast’s 18.4 fps due to its efficient token-wise computation and reduced temporal redundancy. Despite its transformer complexity, TimeSformer maintained lower GPU memory consumption (7.2 GB) compared to SlowFast’s 8.9 GB, attributed to the absence of multi-branch convolutional streams.

From a deployment perspective, both models meet real-time processing thresholds for 25 fps video feeds ^35^. However, the TimeSformer’s higher parallelization efficiency and consistent batch-to-batch latency (∼ 44 ms/frame) make it preferable for continuous inference within digital-twin pipelines where computational scalability and stability are essential. A comparative hardware analysis indicates that real-time deployment is feasible on high-end consumer GPUs or edge-AI modules (e.g., Jetson AGX Orin) with slight down-sampling. Such resource profiling ensures that behavior recognition remains sustainable for on-farm automation without excessive power draw.

#### 3.5.2 Scalability and Multi-Cow Processing

To emulate barn-scale conditions, the end-to-end system was evaluated on simultaneous video streams tracking 16 individual cows using YOLOv11 + ByteTrack for detection and identity assignment. The combined detection-tracking-classification pipeline achieved an average latency of 180 ms per frame, corresponding to real-time throughput at 5.5 fps per cow. Batch-processing optimization and asynchronous GPU queues reduced total inference time by 27 %, confirming the framework’s scalability to group monitoring without significant degradation in accuracy. These benchmarks demonstrate the system’s capacity for large-herd digital-twin synchronization, where multiple physical cows can be updated concurrently in the virtual environment. Future work will explore multi-GPU distribution and temporal batching to achieve near-real-time herd-level behavior analytics.

### 3.6 Challenge Analysis and Performance Limitations

#### 3.6.1 Environmental Variability Impact

Despite strong baseline accuracy, environmental factors continued to influence detection reliability. Under night-time illumination, accuracy declined by 8-12 %, primarily due to reduced contrast and color-channel noise in infrared footage ^9^. Occlusions-caused by barn structures, equipment, or cow overlapped to intermittent trajectory loss, occasionally truncating behavior sequences ^3^. The tracking system successfully reidentified 92 % of interrupted tracks, but brief identity swaps introduced minor temporal labeling noise. Corner and edge regions of the camera field exhibited degraded detection consistency, as cows partially exited the frame, limiting spatial continuity for both TimeSformer and SlowFast. Incorporating multi-view camera fusion and adaptive brightness equalization could mitigate these edge-case degradations.

#### 3.6.2 Behavior-Specific Recognition Challenges

Persistent confusion between Lying and Ruminating & Lying remains the principal classification bottleneck. Both behaviors share near-identical postural geometry; differentiation relies solely on subtle mandibular motion, which is occasionally obscured or temporally aliased at low frame rates ^36^. Enhancing temporal resolution or integrating optical-flow-based motion cues could improve distinction in future iterations. Drinking behavior detection also posed challenges due to reflective water surfaces and variable head angles. False positives occurred when cows lowered their heads near troughs without actual ingestion, emphasizing the need for multi-modal fusion with acoustic or RFID-based water-intake sensors.

Finally, temporal-sequence modeling limitations were evident in transitions between behaviors. Both architectures occasionally produced fragmented predictions during behavior shifts (e.g., standing to lying), suggesting that explicit temporal smoothing or recurrent attention integration could enhance continuity. These findings emphasize that while current deep-video architectures excel at single-state classification, the goal of continuous, biologically faithful behavioral tracking requires hybrid temporal models combining deep attention with probabilistic state transition logic.

## 4. Discussion

The TimeSformer architecture demonstrated distinct advantages for livestock video analytics owing to its global spatiotemporal attention mechanism, which captures both spatial structures and long-term temporal evolution. By dynamically allocating attention to relevant regions such as the head, muzzle, or feed trough, the model maintains interpretability while adapting to motion sparsity typical of commercial barns. This allows robust recognition of prolonged behaviors like ruminating and feeding & standing, whose visual cues evolve slowly over time. In our 24/7 barn recordings, TimeSformer achieved 85.0 % overall accuracy (macro-F1= 0.84) and processed 22.6 fps on an RTX A100, confirming real-time feasibility. The SlowFast architecture, while slightly lower in accuracy (82.3 %), remained competitive for posture-oriented states such as Standing or Lying. Its dual-pathway design (slow spatial, fast motion) efficiently captures short-duration transitions like drinking or standing changes, with predictable latency and low memory overhead, making it attractive for cost-constrained edge deployments. Compared with recent benchmarks CBVD-5 ^37^(687 segments, 107 Holsteins; ∼ 78.7 % accuracy) and the beef-cattle dataset of Cao et al. ^10^ (4974 clips; ∼ 90 % mAP₅₀) our dataset of 4964 annotated clips (expanded to 9600 via augmentation) encompasses seven behavior classes under natural lighting and occlusion, achieving parity with state-of-the-art accuracy despite far noisier conditions. The scale, behavioral granularity, and continuous capture across diurnal cycles make it one of the largest open dairy-behavior video resources, bridging the gap between controlled research corpora and operational barns.

Nonetheless, several methodological limitations merit discussion. Identity leakage remains a primary risk if clips from the same cow appear in both training and test sets, apparent performance may be inflated. Future work will adopt leave-cow-out or day-blocked splits to ensure disjoint identities and report corresponding performance changes. Temporal sampling is another constraint 12 frames per 10-s clip (∼1.2 fps) may undersample mandibular cycles critical for rumination and drinking, ablations at 16–32 frames will clarify the accuracy-latency trade-off. Tracking reliability also warrants quantification, our YOLOv11 and ByteTrack pipeline preserves per-cow IDs but has yet to be evaluated on standard multi-object tracking metrics (MOTA, IDF1, ID-switches) using an annotated subset. Environmental variability further affects performance; accuracy drops 8–12 % at night or under severe occlusion, so stratified tables by day/night, view angle, and occlusion level will be added. Because augmented samples were confined to training, evaluation will be redone on unaugmented test data with natural class priors and balanced accuracy reporting to prevent optimistic bias. The behavioral taxonomy single-label composites such as “lying & feeding” or “standing & ruminating” simplifies ethograms that are inherently multi-label. These were chosen for practical monitoring in tie-stall barns, yet future work will re-analyze behavior along orthogonal posture and include confusion plots separating posture from oral-activity errors.

The structured behavioral outputs cow ID, start/end times, durations, and confidence scores form the behavioral layer of the dairy digital twin. These CSV streams can directly feed NRC feed-intake models to infer dry-matter intake and dynamically adjust rations. Integration within Unity 3D enables real-time visualization of individual cows’ states synchronized with nutritional and environmental modules, establishing an operational link between perception and physiology. Within the twin dashboard, deviations from baseline feeding or rumination cycles act as early-warning indicators of metabolic or welfare issues, supporting proactive interventions. The architecture will be interoperable with commercial herd-management systems such as DairyComp 305 and Lely T4C and will be designed for modular deployment using networked edge devices. Both TimeSformer and SlowFast achieve real-time throughput on mid-range GPUs, ensuring cost-effective scalability. Economic analyses should consider reductions in manual observation labor, improved feed efficiency, and health-related cost savings. Reliability under barn humidity and dust will be maintained through ruggedized enclosures and modular camera design.

Looking ahead, further improvements will emphasize multi-modal fusion, model calibration, and edge optimization. Acoustic and thermal sensors can complement visual data by capturing rumination chewing sounds and heat signatures, improving discrimination under occlusion or darkness. Lightweight transformer variants using pruning, quantization, and adaptive attention weighting will reduce inference cost and enhance interpretability. Federated-learning frameworks can enable cross-farm model updates without sharing raw video, preserving data privacy while addressing regional variability. Quantitative attention-map analyses reporting the proportion of attention mass within anatomical regions of interest will strengthen biological plausibility. Ultimately, integrating behavioral analytics with NRC feed-intake predictions, health metrics, and environmental sensing will create a self-updating digital-twin ecosystem that links perception, prediction, and management. By demonstrating continuous, per-cow behavioral timelines that synchronize physical animals with their virtual counterparts, this framework establishes a scalable foundation for individualized nutrition, early-warning health systems, and climate-smart, welfare-oriented dairy management.

## 5. Data Availability

The annotated video dataset (4,964 clips) and augmentation recipes used to create the 9,600-sample training set include representative sample clips are available on request. Due to facility restrictions, full raw CCTV footage is not publicly released; additional data are available from the corresponding author upon reasonable request.

## 6. Code Availability

All scripts for preprocessing, YOLOv11 detection, ByteTrack tracking, and SlowFast/TimeSformer training and inference are available at https://github.com/mooanalytica/digital-twin-dairycow under the MIT license, with a frozen environment file and instructions to reproduce all reported results.

## 7. Author contributions

S.R. and E.G. curated data, implemented models, performed experiments, and wrote the manuscript. SR contributed to methodology and analysis. SN supervised the project, provided resources and critical revisions. All authors approved the final version.

## 10. Ethics statement

All animal procedures were approved by the Dalhousie University Animal Care and Use Committee (Protocol #2024-026, approval date 16 May 2024) in accordance with Canadian Council on Animal Care (CCAC) guidelines.

## 11. Acknowledgments

The authors gratefully acknowledge the staff and animal caretakers of the Ruminant Animal Centre at Dalhousie University for their assistance with data collection.

